# Streamlined Structure Determination by Cryo-Electron Tomography and Subtomogram Averaging using TomoBEAR

**DOI:** 10.1101/2023.01.10.523437

**Authors:** Nikita Balyschew, Artsemi Yushkevich, Vasilii Mikirtumov, Ricardo M. Sanchez, Thiemo Sprink, Misha Kudryashev

**Affiliations:** Max Planck Institute of Biophysics, Frankfurt on Main, Germany; Buchmann Institute for Molecular Life Sciences, Goethe University of Frankfurt on Main, Germany; *In Situ* Structural Biology, Max Delbrück Centre for Molecular Medicine in the Helmholtz Association, Berlin, Germany; Core Facility for Cryo-Electron Microscopy, Charité-Universitätsmedizin Berlin, Germany; Institute of Medical Physics and Biophysics, Charité-Universitätsmedizin Berlin, Germany; Cryo-EM Facility, Max Delbrück Centre for Molecular Medicine in the Helmholtz Association, Berlin, Germany; EMBL Heidelberg, Germany

## Abstract

Structures of macromolecules in their native state provide unique unambiguous insights into their functions. Cryo-electron tomography combined with subtomogram averaging demonstrated the power to solve such structures *in situ* at resolutions in the range of 3 Angstrom for some macromolecules. In order to be applicable to structural determination of the majority of macromolecules observable in cells in limited amounts, processing of tomographic data has to be performed in a high-throughput manner. Here we present TomoBEAR - a modular configurable workflow engine for streamlined processing of cryo-electron tomographic data for subtomogram averaging. TomoBEAR combines commonly used cryo-EM packages and reasonable presets to provide a transparent “white box” for data management and processing. We demonstrate applications of TomoBEAR to two datasets of purified proteins and to a membrane protein RyR1 in a membrane and demonstrate the ability to produce high resolution with minimal human intervention. TomoBEAR is an open-source and extendable package, it will accelerate the adoption of *in situ* structural biology by cryo-ET.

## INTRODUCTION

Cryo-electron tomography (cryo-ET) enables observation of natively preserved molecules in the context of intact cells (Beck & Baumeister, 2016). Combination of cryo-ET with subtomogram averaging (StA) allows obtaining angstrom-scale structures of macromolecules (Wan & Briggs, 2016; Leigh *et al*, 2019). StA provided unique insights into structure and function of viral and bacterial structural proteins (Schur *et al*, 2016; von Kügelgen *et al*, 2020), eukaryotic protein coats (Hutchings *et al*, 2021), actin filaments in sarcomeres (Wang *et al*, 2022) and even visualizing ribosomes inside intact bacterial cells (Xue *et al*, 2022) with bound small molecules (Tegunov *et al*, 2021). Developments in hardware and software (Mastronarde, 2005; Mastronarde & Held, 2017; Castaño-Díez *et al*, 2012, 2016; Jiménez de la Morena *et al*, 2022; Turoňová *et al*, 2017) allow obtaining higher resolution for the broader range of samples. In particular, recently several approaches allowed refinement of non-linear sample movement and of electron optical distortions allowing significant resolution improvement (Himes & Zhang,2018; Chen *et al*, 2019; Tegunov *et al*, 2021; Zivanov *et al*, 2022).

However, several major hurdles remain in making StA a mainstream method. First – the StA workflow includes multiple steps that are typically performed by specialized software packages (Leigh *et al*, 2019) which require special effort to interface (Burt *et al*, 2021). Second – several steps in the StA workflow – tilt-series alignment and particle identification often require manual intervention. Third – optimal storing and processing of large amounts of 3D volumes requires large-scale computing infrastructure and is not straightforward even for expert users. Large amounts of intermediate results such as tomograms, extracted particles, metadata may occupy terabytes of hard drive space and need to be managed. Finally, most macromolecules occur in cells in limited copy numbers therefore in order to achieve meaningful resolution, it is necessary to record and process large amounts of data. Excitingly, there is significant progress in speeding up tomographic data collection by recording tilt-series in parallel (Bouvette *et al*, 2021; Eisenstein *et al*, 2022). In order to process large amounts of data, the processing software also needs to be designed for automation, yet with opportunities for user interventions where it is needed.

Here we present TomoBEAR (Basics of Electron tomography and Automatic Reconstruction) which is an open-source workflow for at-scale mostly automated pipeline for structure determination from cryo-electron tomograms. TomoBEAR interfaces commonly used software for cryo-ET and allows flexibility for users to develop pipelines for their molecules of interest. We demonstrate applications of TomoBEAR to three datasets reaching high resolution with minimal user input.

## RESULTS

### Overall design and processing options

TomoBEAR is implemented as a modular pipeline runner, executing one module per one tilt-stack, a tomogram or a set of particles. TomoBEAR can run in parallel in high-performance computing environments such as multi-GPU workstations and/or HPC clusters. TomoBEAR takes the output of data collection - movie frames stored on the hard drive as input, generates metadata and performs motion correction with MotionCorr2 (Zheng *et al*, 2017), assembly of tilt-series, tilt-series alignment with IMOD (Mastronarde & Held, 2017), Dynamo (Castaño-Díez *et al*, 2012) or AreTomo (Zheng *et al*, 2022), defocus determination with GCTF (Zhang, 2016) and 2D correction of contrast transfer function (CTF) followed by tomographic reconstruction using IMOD (Kremer *et al*, 1996). Up to this step the workflow operates in a near-automated manner and could be used for live data processing during data collection (Figure 1A).

**Figure 1.**
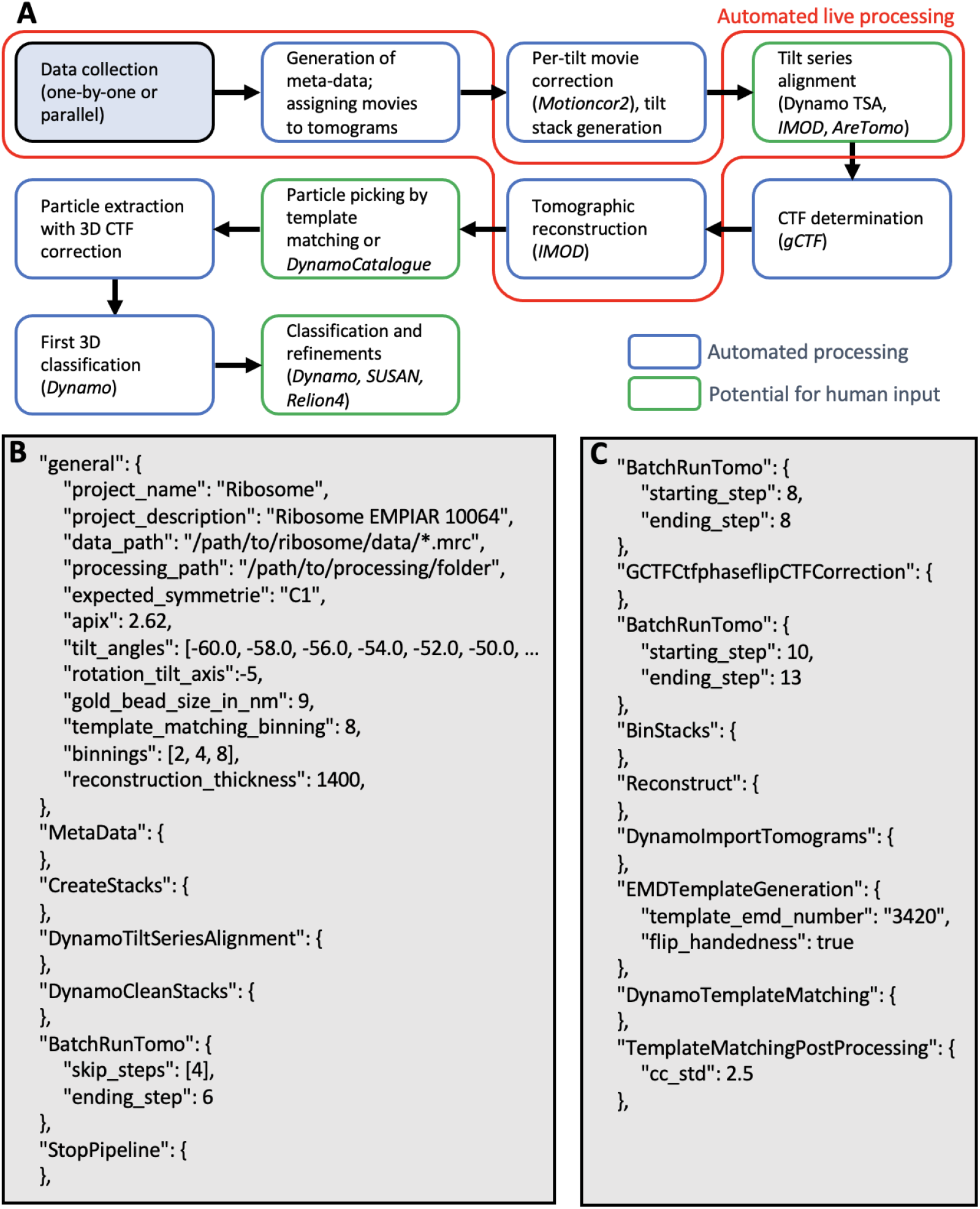
A standard workflow for data processing in TomoBEAR. A) A flow diagram of data processing with TomoBEAR. Blue boxes outline the steps that are performed fully automatically, green boxes may require human intervention. The steps highlighted in red highlight the functionality of live data processing. B,C) an example of a .json configuration file that was used for processing of the EMPIAR-10064 dataset (results below). Panel B contains the general module and the steps for processing up to refinement of gold fiducials. Panel C contains modules for CTF correction, tomographic reconstruction and template matching.

For particle picking and subtomogram averaging several options are available depending on target molecules. If the molecules of interest can be identified in tomograms by template matching, it can be done automatically followed by particle extraction to the hard-drive, generation and execution of the Dynamo-based classification project for initial subtomogram classification by multi-reference alignment using Dynamo (Castaño-Díez *et al*, 2012). Further alignment projects can be generated by selecting classes; upon changes in the desired binning levels - a selection of particles can be generated by direct reconstruction from tilt-series, without the need to reconstruct large unbinned tomograms.

When the molecules of interest can not be picked by template matching - TomoBEAR creates data structures for the Dynamo catalogue system (Castaño-Díez *et al*, 2016) - a previously reported versatile tool for geometry-supported particle picking. For high-resolution structure determination metadata can be exported to Relion (Zivanov *et al*, 2022) or the substacking analysis tool SUSAN (https://github.com/KudryashevLab/SUSAN, manuscript in preparation) based on projective cross-correlation (Sánchez *et al*, 2019b, 2019a).

TomoBEAR implements a “white box”approach that allows users to keep track of the used parameters and to monitor intermediate results. Modules can be re-run for the entire dataset or for selected tilt-stacks/tomograms providing an opportunity to test the pipeline on a subset of data to tune parameters and then process the entire dataset.

### Automated tilt-series preprocessing and tomographic reconstruction

The user can define the execution parameters in a *.json* file (example in Figure 1B,C Supplementary Text 2) which contains a *general* section describing the overall parameters and module-specific sections with module-specific parameters. A set of default parameters is provided in *defaults.json*, users can modify them by passing the parameters to the module description. Upon execution the TomoBEAR output folder is populated by module-specific folders. Typically each operation creates a sub-folder for each tilt-series/tomogram that can be inspected and upon a successful completion a SUCCESS file is created inside the corresponding folder. Upon a successful execution of the module for all the tomograms a global SUCCESS file is written into the module-specific folder. In order to re-execute the module the global SUCCESS file can be deleted and the execution can be re-started.

The first operations of the cryo-ET and subtomogram averaging pipeline implemented in TomoBEAR are performed automatically. An output folder from a microscopy session (original movie frames) can be the input for TomoBEAR. Files are assigned to tilt-series (*SortFiles* module), drift is corrected (*MotionCorr2* module) and the IMOD-compatible stacks are assembled (*CreateStacks* module). Batch-alignment of tilt-series may be performed using gold fiducials with IMOD’s BatchRunTomo (Mastronarde & Held, 2017) or Dynamo (Castaño-Díez *et al*, 2012), or using patch tracking in IMOD or AreTomo (Zheng *et al*, 2022). The latter option is useful for aligning tilt-series from cryo FIB-milled lamellae. We found tilt-series alignment with the recently released tilt-series alignment algorithm DynamoTSA developed by Daniel Castano-Diez (https://wiki.dynamo.biozentrum.unibas.ch/w/index.php/Walkthrough_on_GUI_based_tilt_series_alignment) to be robust, precise and well performing computationally, therefore TomoBEAR uses it as default when gold fiducials are available. Importantly, DynamoTSA contains a robust estimation of whether the alignment was successful or not; this check allows the users to selectively inspect failed tilt-series. The resulting fiducial-based or fiducial-less alignments optionally can be inspected and potentially refined in a parallelly maintained IMOD project. The module *BatchRunTomo* allows execution of the steps as they are defined in IMOD (Mastronarde & Held, 2017) (Supplementary Text 1); TomoBEAR uses the IMOD project for calculating and optionally refining the final tomographic alignment parameters, CTF correction and tomographic reconstruction. Initial defocus determination and 2D CTF correction is performed automatically using gCTF (Zhang, 2016) and *Ctfphasefilp* from IMOD (Xiong *et al*, 2009) in the *GCTFCtfphaseflipCTFCorrection* module. The fits of experimental and estimated power spectra can be examined in the processing folder. Aligned, binned, CTF corrected tilt-stacks are produced and the tomograms are reconstructed at the user-defined binning levels.

### Particle picking

Large proteins and lattices can be successfully picked by template matching (Frangakis *et al*, 2002). In TomoBEAR we reimplemented the Dynamo (Castaño-Díez *et al*, 2012) version of template matching on graphical processors, speeding up the processing 12-15 times and developed auxiliary tools. A template for the search can be provided by a user as a file or it can be automatically produced from an EMD entry by resampling to the voxel size of a tomogram and low-pass filtering it to conservative resolution. The cross-correlation map resulting from template matching can be post-processed, such as removing “large islands’’ and/or connected regions like membranes/edges (*TemplateMatchingPostProcessing* module). The user can inspect the cross-correlation (CC) maps and if the peaks are well-defined at the positions of the proteins of interest - particle extraction can be performed. For this the positions of the top hits are extracted to a Dynamo-style table limited by threshold criteria (standard deviations compared to the mean CC value and/or maximal number of particles per tomogram; *TemplateMatchingPostProcessing* module). Dynamo-style table in this case contains initial orientations of particles that can be used as an input for local subtomogram alignment, classification and averaging.

When particles cannot be picked by template matching – a versatile set of particle-picking tools is available in the previously described Dynamo catalogue system (Castaño-Díez *et al*, 2016) which can be created with the *DynamoImportTomograms* module. Particle picking is generally performed in highly binned tomograms which can potentially be filtered (Frangakis, 2021) for visualization purposes. As a result of particle picking/template matching the initial subtomograms are either extracted from existing tomograms or are reconstructed from tilt-series at the defined binning levels. For particles reconstructed from projections, 3D CTF correction is performed taking the height of the particle in the tomogram into account for CTF correction. Particle reconstruction from projection is performed on GPUs using the SUSAN engine (https://github.com/KudryashevLab/SUSAN). Additionally, to a significant speedup in the execution time, reconstruction of particles from projections eliminates the need to reconstruct large unbinned tomograms, saving hard drive space.

### Subtomogram alignment, classification and averaging

A multi-reference alignment and classification project with Dynamo can be generated and executed automatically (*DynamoAlignmentProject* module) with pre-calculated parameters. In our implementation we use the originally used templates as well as “noise traps” as references for multi-reference alignment. This allows to separate true/good particles and false positives/bad particles obtained by template matching or semi-automated particle picking. Consecutive Dynamo classification projects may be initiated after class selection by the user or several classification steps can be scheduled at the start of the processing (see example in Supplementary Text 2). Binning can be reduced, in this case a new set of particles will be produced either by cropping particles by reconstructed tomograms at lower binning or by direct reconstruction (“SUSAN particles”). At each step of particle extraction, the particles are recentered. When a final particle set is produced, independent half-set refinement is performed in order to reliably assess the resolution. We call this processing workflow “conventional StA”.

Generally, the automation of StA at the final steps of the refinement has limited utility as the user has a lot of flexibility to optimize particle sets, masks, filters and other parameters. We tested the workflow for several datasets (below) and found that for simple objects with a well-tailored configuration, near-automated StA can reveal results close to optimal. Ultimately, Dynamo and similar approaches operating on 3D volumes that contain misalignments, have limitations in obtainable resolution. Therefore, we implemented export to SUSAN (https://github.com/KudryashevLab/SUSAN) by creating the data structures and to Relion (Zivanov *et al*, 2022) by creating *.star* files with tomograms and particle descriptions. The mentioned *.star* file can also be used for processing hybridStA data (Sanchez *et al*, 2020; Song *et al*, 2020) with other single particle packages such as CryoSparc (Punjani *et al*, 2017).

### TomoBEAR utilities

Some operations of the cryo-ET/subtomogram averaging workflow require human intervention. As mentioned, TomoBEAR maintains a parallel IMOD project during tilt-series processing, which provides compatibility with IMOD’s wide functionality. That allows visual inspection of the processing steps, and, importantly, inspection and refinement of the gold fiducial-based or fiducial-less tilt-series alignment. A module *StopPipeline* can be added to the configuration file to stop automated execution for these purposes.

We realized that often fiducial gold beads differ in sizes and specifying variable sizes helps automatic algorithms to succeed in tilt-series alignment. Therefore, multiple gold bead sizes may be input to the *general* module. In this case tilt-series alignment routine will try all the sizes consecutively till the first successful alignment.

Note that in case of particle picking by template matching, false positive picks may be produced by gold beads or their shadows, edges of grid holes, edge artifacts from reconstructions, contaminations on ice surfaces and other sources. TomoBEAR aims to minimize the amount of such false positives by erasing gold beads by default (IMOD implementation in 2D), smoothing of image edges (as in Figure 2A) and an option to erase grid edges from micrographs (*GridEdgeEraser* module).

**Figure 2.**
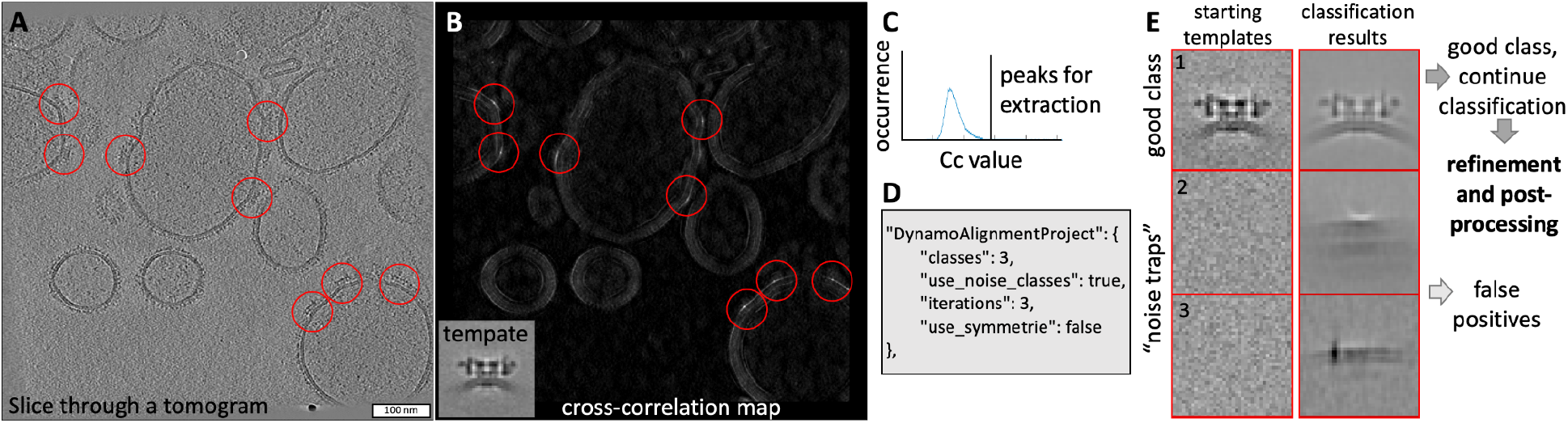
Example of particle picking and subtomogram classification from a tomogram of an ion channel RyR1 imaged in native membranes (EMPIAR-10452). A) A slice through a tomogram with particles of RyR1 marked with red circles. B) A result of the Dynamo-style cross-correlation search of a low-resolution template (shown in an inset) in a tomogram from (A) with the 15 degree angular search and C4 symmetry. C) Selecting an offset for the CC-values of the extracted peaks based on the histogram of the CC-values in the map in (B). D) A TomoBEAR description of a classification project to run multi reference alignment with 3 classes, two of which are noise traps for 3 iterations. E) Classification of particles extracted from the positions corresponding to the top values of the cross-correlation map starting with 3 classes, two of which are noise classes. Resulting class one contains particles which can be further processed, classes 2 and 3 - false positives.

TomoBEAR allows producing tomographic reconstructions “live” in order to probe the quality of the output data during data collection. For simplicity and speed-up TomoBEAR-live skips motion and CTF correction and performs reconstruction on binned input stacks. For initialization users need to provide a *.json* file with the conventional set of parameters plus the expected number of images per tilt-series and the listening time threshold. We tested live data processing to generate tomographic reconstructions of the previously reported HIV-1 GAG dataset (Schur *et al*, 2016), EMPIAR 10164. For the tests we simulated live data collection with a script that copied original data to the simulated data collection folder updating original timestamps. Frame-to-frame and tilt-serie to tilt-serie time delays were set to 5 and 30 seconds imitating data collection. We measured times spent by TomoBEAR to process data from raw dose-fractionated movies up to reconstructed tomograms in both conventional (offline) and live modes (Figure S1). The simplified workflow aimed at visualization of tomograms resulted in a 3-fold speedup for processing data in live mode compared to the conventional one.

Data management and cleanup: as stand-alone packages, many single particle, tomography and StA utilities produce temporary files that can be removed using the cleanup functionality of TomoBEAR during or after the execution.

### Benchmarking TomoBEAR

We benchmarked TomoBEAR on two previously reported and one original dataset, the overview of processing time and results can be found in Table 1. First, we processed the tomograms of purified 80S ribosomes imaged by cryo-ET (Khoshouei *et al*, 2017, EMPIAR 10064, mixedCTEM). This dataset was already motion-corrected and tilt-series assembled. We performed automated conventional data processing with particle picking by template matching and subtomogram classification and averaging in Dynamo. The user inputs were: minor refinements of gold fiducial alignment and class selection during subtomogram classification with Dynamo. The resulting resolution was 11.3 Å (Figure 3A), similar to the 11.2 Å resolution reported in the original publication. The processing time was ~1h for preprocessing from assembled and motion-corrected stacks up to reconstruction and ~13h 30m for template matching and StA analysis.

**Table 1.**
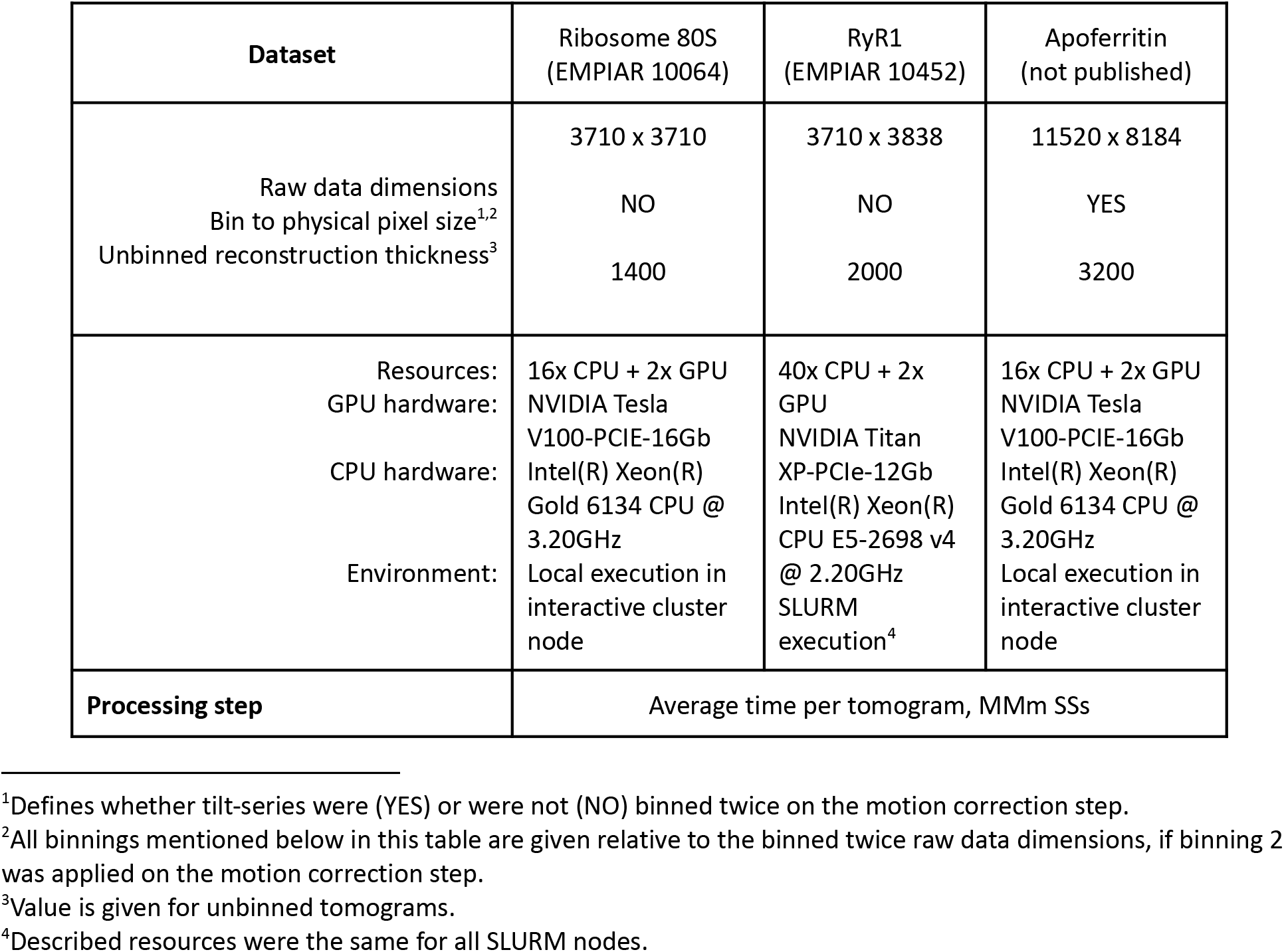
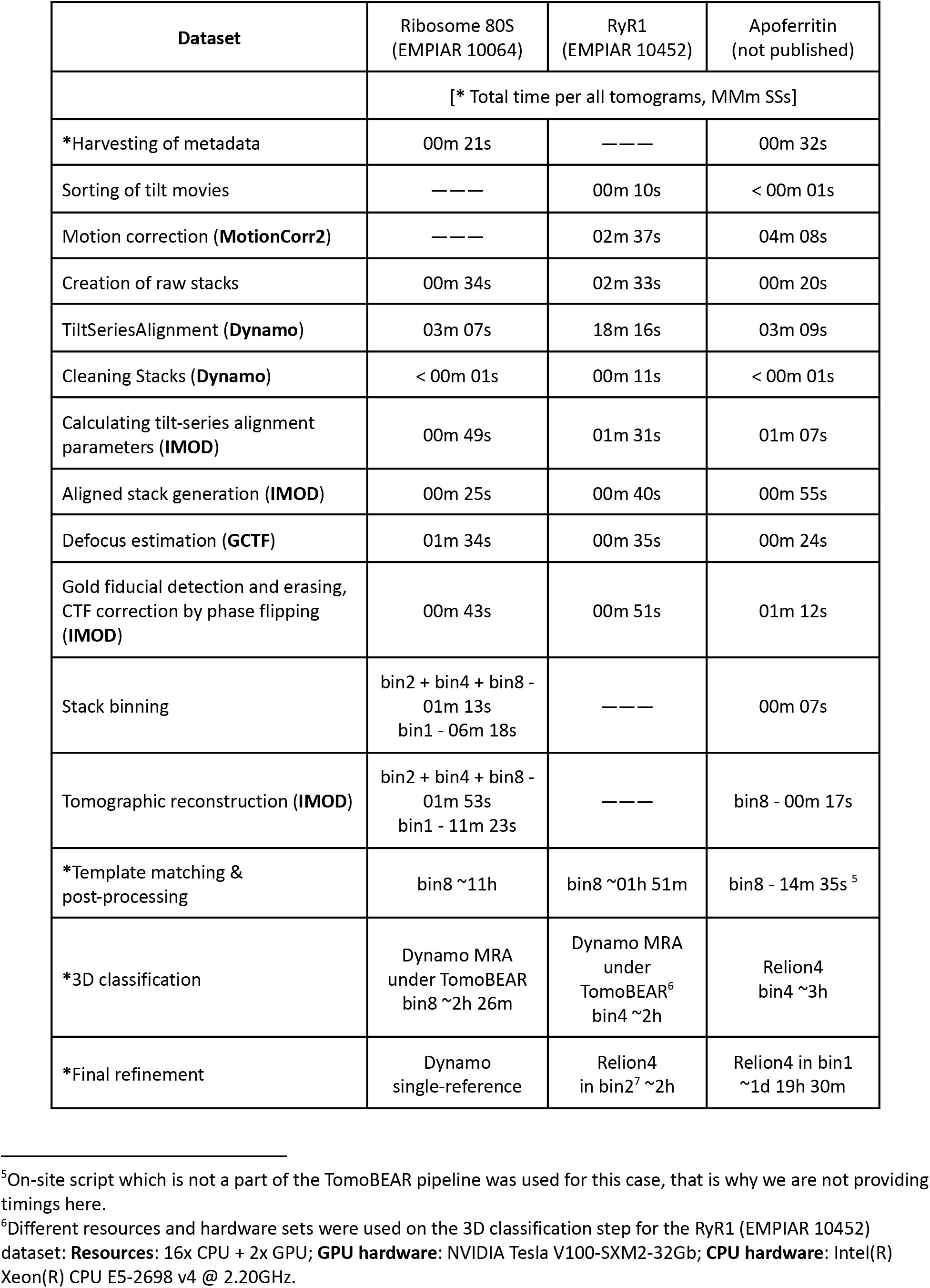
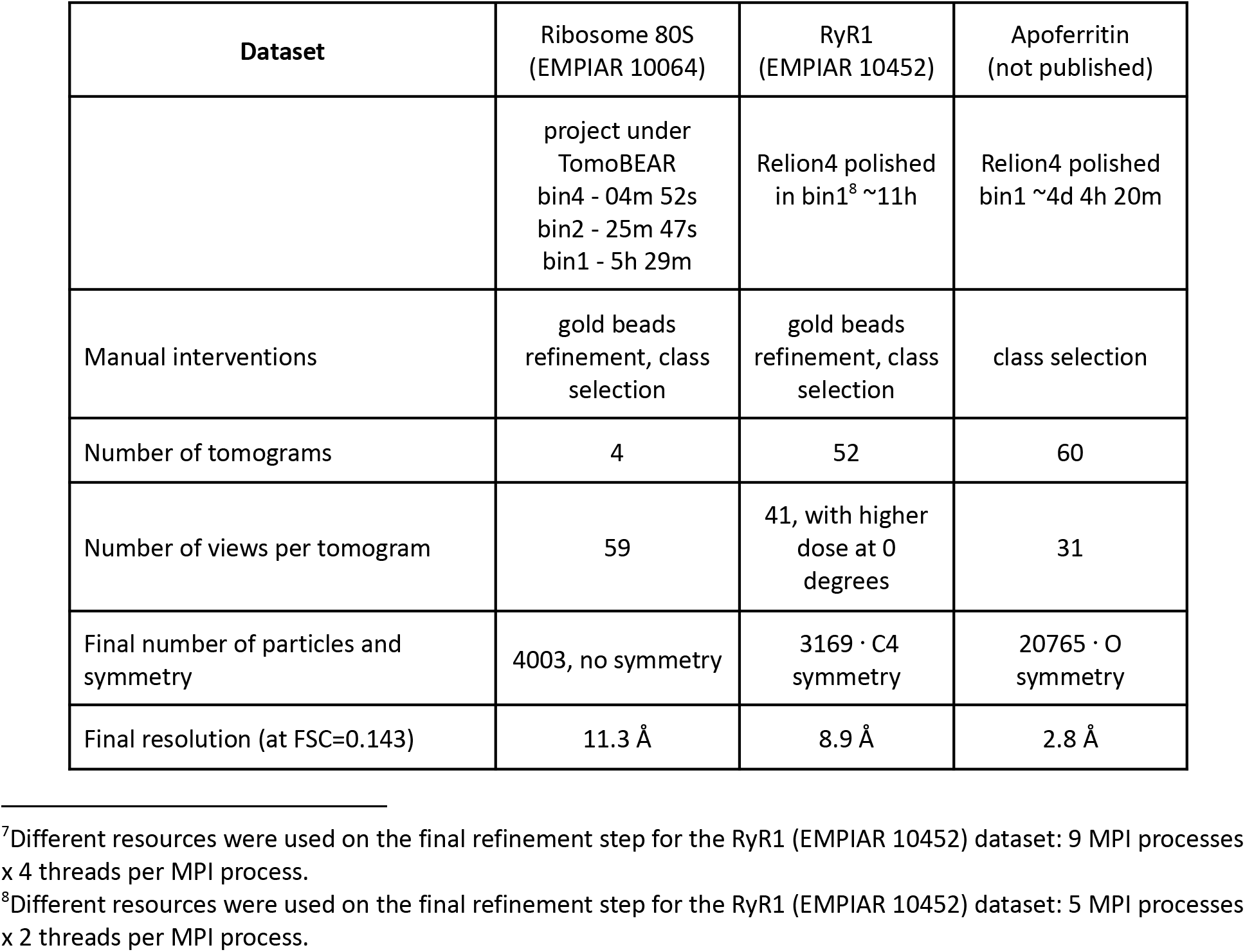
Summary for processing of the benchmarking datasets.

| Dataset | Ribosome 80S<br>(EMPIAR 10064) | RyR1<br>(EMPIAR 10452) | Apo ferritin<br>(not published) |
| --- | --- | --- | --- |
| Raw data dimensions | 3710 x 3710 | 3710 x 3838 | 11520 x 8184 |
| Bin to physical pixel size<sup>1,2</sup> | NO | NO | YES |
| Unbinned reconstruction thickness<sup>3</sup> | 1400 | 2000 | 3200 |
| Resources: | 16x CPU + 2x GPU | 40x CPU + 2x GPU | 16x CPU + 2x GPU |
| GPU hardware: | NVIDIA Tesla V100-PCIe-16Gb | NVIDIA Titan XP-PCIe-12Gb | NVIDIA Tesla V100-PCIe-16Gb |
| CPU hardware: | Intel(R) Xeon(R) Gold 6134 CPU @ 3.20GHz | Intel(R) Xeon(R) CPU E5-2698 v4 @ 2.20GHz | Intel(R) Xeon(R) Gold 6134 CPU @ 3.20GHz |
| Environment: | Local execution in interactive cluster node | SLURM execution<sup>4</sup> | Local execution in interactive cluster node |
| Processing step | Average time per tomogram, MMm SSs |  |  |

<sup>1</sup>Defines whether tilt-series were (YES) or were not (NO) binned twice on the motion correction step.
<sup>2</sup>All binnings mentioned below in this table are given relative to the binned twice raw data dimensions, if binning 2 was applied on the motion correction step.
<sup>3</sup>Value is given for unbinned tomograms.
<sup>4</sup>Described resources were the same for all SLURM nodes.

| Dataset | Ribosome 80S<br>(EMPIAR 10064) | RyR1<br>(EMPIAR 10452) | Apoferritin<br>(not published) |
| --- | --- | --- | --- |
|  | [* Total time per all tomograms, MMm SSs] |  |  |
| *Harvesting of metadata | 00m 21s | --- | 00m 32s |
| Sorting of tilt movies | --- | 00m 10s | < 00m 01s |
| Motion correction (**MotionCorr2**) | --- | 02m 37s | 04m 08s |
| Creation of raw stacks | 00m 34s | 02m 33s | 00m 20s |
| TiltSeriesAlignment (**Dynamo**) | 03m 07s | 18m 16s | 03m 09s |
| Cleaning Stacks (**Dynamo**) | < 00m 01s | 00m 11s | < 00m 01s |
| Calculating tilt-series alignment parameters (**IMOD**) | 00m 49s | 01m 31s | 01m 07s |
| Aligned stack generation (**IMOD**) | 00m 25s | 00m 40s | 00m 55s |
| Defocus estimation (**GCTF**) | 01m 34s | 00m 35s | 00m 24s |
| Gold fiducial detection and erasing, CTF correction by phase flipping (**IMOD**) | 00m 43s | 00m 51s | 01m 12s |
| Stack binning | bin2 + bin4 + bin8 -<br>01m 13s<br>bin1 - 06m 18s | --- | 00m 07s |
| Tomographic reconstruction (**IMOD**) | bin2 + bin4 + bin8 -<br>01m 53s<br>bin1 - 11m 23s | --- | bin8 - 00m 17s |
| *Template matching & post-processing | bin8 ~11h | bin8 ~01h 51m | bin8 - 14m 35s <sup>5</sup> |
| *3D classification | Dynamo MRA<br>under TomoBEAR<br>bin8 ~2h 26m | Dynamo MRA<br>under<br>TomoBEAR<sup>6</sup><br>bin4 ~2h | Relion4<br>bin4 ~3h |
| *Final refinement | Dynamo<br>single-reference | Relion4<br>in bin2<sup>7</sup> ~2h | Relion4 in bin1<br>~1d 19h 30m |

<sup>5</sup>On-site script which is not a part of the TomoBEAR pipeline was used for this case, that is why we are not providing timings here.
<sup>6</sup>Different resources and hardware sets were used on the 3D classification step for the RyR1 (EMPIAR 10452) dataset: **Resources:** 16x CPU + 2x GPU; **GPU hardware:** NVIDIA Tesla V100-SXM2-32Gb; **CPU hardware:** Intel(R) Xeon(R) CPU E5-2698 v4 @ 2.20GHz.

| Dataset | Ribosome 80S<br>(EMPIAR 10064) | RyR1<br>(EMPIAR 10452) | Apoferitin<br>(not published) |
| --- | --- | --- | --- |
|  | project under<br>TomoBEAR<br>bin4 - 04m 52s<br>bin2 - 25m 47s<br>bin1 - 5h 29m | Relion4 polished<br>in bin1<sup>8</sup> ~11h | Relion4 polished<br>bin1 ~4d 4h 20m |
| Manual interventions | gold beads<br>refinement, class<br>selection | gold beads<br>refinement, class<br>selection | class selection |
| Number of tomograms | 4 | 52 | 60 |
| Number of views per tomogram | 59 | 41, with higher<br>dose at 0<br>degrees | 31 |
| Final number of particles and<br>symmetry | 4003, no symmetry | 3169 · C4<br>symmetry | 20765 · O<br>symmetry |
| Final resolution (at FSC=0.143) | 11.3 Å | 8.9 Å | 2.8 Å |

**Figure 3.**
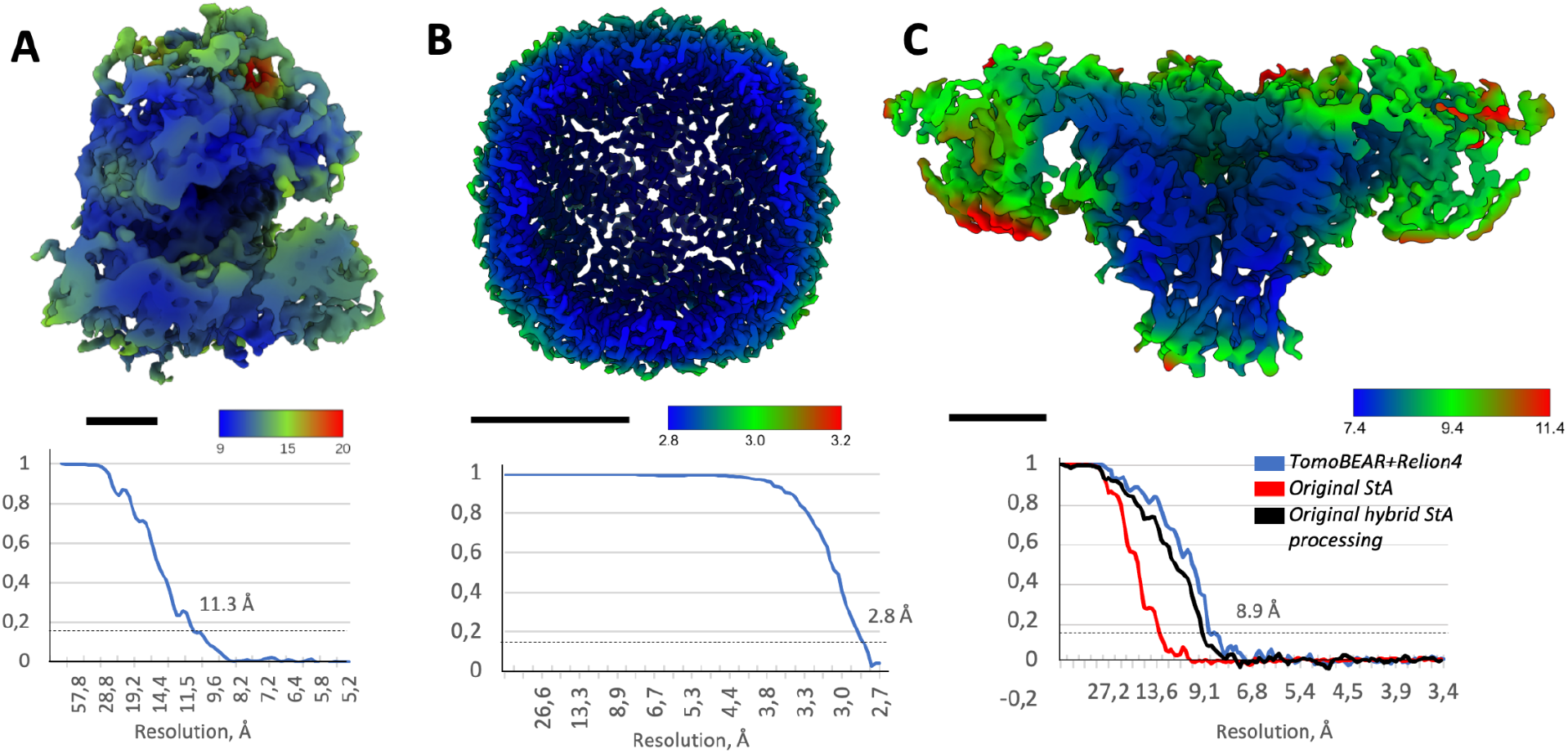
Benchmarking performance of TomoBEAR. A) Processing of the tomographic datasets of purified ribosomes (EMPIAR 10064, mixed defocus): A structure at resolution 11.3 Å sliced in the middle of the reconstruction and colored according to local resolution. Black scale bars: 10 nm. Lower panels: detection of resolution based on Fourier Shell correlation between independently refined half-sets. B) Structure of purified human apoferritin imaged by cryo-ET in this study at an overall resolution of 2.8 Å. The map is sliced in the middle of the reconstruction and colored according to local resolution. C) Processing of tomograms of an ion channel RyR1 imaged in native membranes purified from rabbit muscle (EMPIAR-10452). Structure of RyR1 at a resolution 8.9 Å sliced in the middle of the reconstruction and colored according to local resolution. Estimation of resolution based on Fourier Shell Correlation: the curves for the original processing of this dataset from ref. (Sanchez et al, 2020) are in red and black, blue - from TomoBEAR+Relion4.

Next, we recorded 60 tomograms of purified human apoferritin and performed data processing with TomoBEAR followed by export to Relion4 for StA. In this case we performed tilt-series alignments by TomoBEAR without manual inspection or corrections. Particles were picked by template matching with reduced angular search range due to high symmetry of the target molecule. Particle set was imported to SUSAN and then to Relion4 after the initial alignment. Processing with Relion4 was done manually, two rounds of Refine3D, particle polishing and CTF refinement of 18k particles with O-symmetry in Relion4 resulted in a structure at a resolution of 2.8 Å, almost reaching the Nyquist limit of 2.7 Å (Figure 3B). We didn’t proceed with StA in Relion4 in super-resolution because of high requirements for computational resources (box size 400 px). For this dataset processing with TomoBEAR up to export in Relion was fully automated. The total runtime of TomoBEAR on 60 tomograms with the outlined hardware setup was around 1 week, the details are summarized in Table 1. The time spent on subtomogram averaging consisted mostly of Refine3D runs; the speed limited by the available hardware and a relatively high number of particles needed to reach the resolution close to Nyquist. In addition to that, the jobs spent some time queueing for the cluster resources. Nevertheless, the final map was obtained within 2 weeks from the day when the TomoBEAR processing was finished.

Finally, we re-processed our previously reported dataset of the ion channel RyR1 imaged in native membranes purified from rabbit muscle imaged by hybridStA (Sanchez *et al*, 2020), EMPIAR-10452. We performed automated tilt-series processing with minor refinement of gold fiducials followed by export to Relion4. Final refinement and particle polishing with Relion4 resulted in a structure at 8.9 Å resolution, slightly surpassing the originally reported resolution of 9.1 Å (Figure 3C). Interestingly, the original structure was produced only from the unitilted image recorded with higher dose (15 e^-^/Å ^2^). Here low-tilted images also contributed to the structures, slightly improving the resulting resolution, although the final particle sets were also slightly different to the previously reported. While it’s hard to accurately estimate the processing time spent for the structure in the original publication, it was in the order of months for particle picking, classifications and refinements. In contrast, only 1 round of Dynamo multireference alignment in the TomoBEAR environment was enough to obtain a homogeneous particle set suitable for the downstream refinement. This allowed us to perform the subtomogram averaging and obtain the final map within one week, thus dramatically reducing the total processing time compared to the original mostly manual processing (Chen & Kudryashev, 2020; Sanchez *et al*, 2020).

## DISCUSSION

Here we report the workflow that enables at-scale processing of cryo-ET data for subtomogram averaging, which is becoming increasingly necessary. Most macromolecules are present in cells in limited copy numbers (Wiśniewski *et al*, 2014); determining their structures *in situ* is only possible by processing large numbers of particles for alignment and averaging. While overexpression can be helpful in some cases, recording a high number of tilt-series is a generally applicable solution. Cryo-ET technology becomes more accessible due to larger microscope install base and the increasingly popular application of the “synchrotron model” (Walsh *et al*,2022). Furthermore, the throughput of data collection dramatically increases by recording tomograms in parallel (Bouvette *et al*, 2021; Eisenstein *et al*, 2022). Therefore, recording large datasets becomes less critical than processing them. Manual processing requires time and expert knowledge of software and/or scripting and has risks of errors and inconsistencies. Suboptimal processing can not only limit the resulting resolution, but also unnecessarily amplify the amount of the used computational resources or of hard drive storage. Therefore the use of streamlining of processing with workflows like TomoBEAR will ease the entry barriers to the StA and will improve the quality of the resulting structures.

Several excellent workflows have been previously released, among others: IMOD+BatchRunTomo (Mastronarde & Held, 2017), emClarity (Himes & Zhang, 2018), tomoAuto (Morado *et al*, 2016), Dynamo (Scaramuzza & Castaño-Díez, 2021), EMAN2 (Chen *et al*, 2019), “M” combined with IMOD and Relion (Tegunov *et al*, 2021), ScipionTomo (Jiménez de la Morena *et al*, 2022). The design of TomoBEAR is aimed at minimizing user intervention for large-scale data processing, focusing on high-resolution StA structures and maintaining flexibility. For this, specific modifications have been implemented such as a good set of default values, minimization of false-positives for template matching, etc. Indeed, we showed that with minimal user interventions it is possible to process mid-scale datasets in a short time and to obtain up-to-date structures. Steps of TomoBEAR scale linearly with the data and can be used for larger datasets, over 500 tomograms (Kudryashev group, unpublished). As TomoBEAR is modular, the addition of further functionality is possible that can be used for optimizing speed and performance of the StA modules. In particular, due to high information content in cryo-electron tomograms, currently particle picking is a challenging step requiring significant computational time or manual work. Neuronal network-based particle pickers (Moebel *et al*,2021; Chen *et al*, 2017) and non-supervised tomogram annotation (Rice *et al*, 2022) tools could be further incorporated to the workflow pre-trained or with pre-defined parameters where possible. Furthermore, a streamlined workflow can be used to systematically evaluate the performance of algorithms for speed and the impact on the final structures. Importantly, in the future automatically processed subset of data can be used as training for neuronal networks that in turn can process large-scale datasets.

Here we presented three applications where minimal user intervention was applied and up-to-date structures were obtained for the benchmarking datasets. For the new dataset of purified human apoferritin we could reach the resolution better than 3 Å, close to the sampling limit, suggesting that the used processing steps do not restrict the obtainable resolution. A dataset or RyR1 contains a membrane protein which are considered to be more difficult targets for particle picking and alignment to the average. Therefore, we believe that our workflow is applicable to a wide range of datasets and can routinely be used to extract maximum information from data. Several recent processing approaches enable the refinement of imperfections accumulated during processing at the final step of structural determination (Himes & Zhang, 2018; Tegunov *et al*, 2021; Chen *et al*, 2019; Zivanov *et al*, 2022). The use of final refinement steps (Relion4 in two of our benchmarking datasets) is therefore highly beneficial for the workflows as it allows more flexibility and a certain margin of error for the processing, making automated processing more robust. Taken together, we believe that the use of TomoBEAR will lower the entry barriers into cryo-ET and subtomogram averaging and will speed up high-resolution structural analysis of macromolecules in their native state.

## MATERIALS AND METHODS

TomoBEAR is a configurable and customizable software package specialized for cryo-electron tomography and subtomogram averaging written in MATLAB (Mathworks). TomoBEAR implements Basics for Electron tomography and Automated Reconstruction of cryo-electron tomography data. TomoBEAR is designed to operate on data in parallel where it is possible and to minimize user interventions and the need to learn the different software packages (MotionCor2, IMOD, Gctf, Dynamo) to be able to process cryo-electron tomography data. TomoBEAR contains a set of predefined defaults stored in the *defaults.json* file, which worked for most of the processing steps that we tested. However, if some parameters need to be modified for specific projects, the parameters can be further customized in a *.json* configuration file. The description of the modules is provided in Supplementary Text 1; Supplementary Text 2 contains a workflow for structural determination of the ribosome benchmarking dataset.

### Data processing details

#### 1. 80S ribosomes (EMPIAR-10064, MixedCTEM)

This is a previously reported conventional transmission electron microscopy (CTEM) dataset with mixed defocus values from the Volta phase plate report (Khoshouei *et al*, 2017). The raw tilt movies were already gain- and motion corrected, and assembled into 4 tilt stacks which were used as an input data for TomoBEAR. Further automated conventional data processing was performed for each step unless otherwise stated. Input tilt stacks were subjected to Dynamo tilt-series alignment routines to produce the fiducial alignment models that were imported to ETOMO from IMOD (Kremer *et al*, 1996) and inspected manually, followed by generation of the stacks. GCTF (Zhang, 2016) was used to estimate defocus values for each tilt, and these values were input to *ctfphaseflip* (Xiong *et al*, 2009) of ETOMO, followed by erasing of gold fiducials. CTF-corrected gold-erased aligned stacks were binned by a factor of 8, bin8 tomograms were reconstructed and subjected to template matching, with the first template being a low-pass-filtered map of the 80S ribosome from this dataset (EMD-3420). 4490 top-hits were extracted and subjected to several subsequent multi-reference alignment projects in Dynamo each with 4 classes. Afterwards, the best class containing 4005 particles was selected manually after visual inspection. Subsequent subtomogram classification and averaging was performed automatically using Dynamo within TomoBEAR. Particles for the final refinement were produced by direct reconstruction by SUSAN refinement for unbinned particles and resulted in 4003 particles averaged to produce the final map. The global resolution of the final map was estimated with 0.143 criterion to be 11.3 Å, similar to the previously reported 11.2 Å with this data in the original publication (Khoshouei *et al*, 2017).

#### 2. Purified human apoferritin

Human apoferritin was produced according to the Leicester protocol (Sheng *et al*, 2022) and diluted with a buffer containing 5 nm colloidal gold to a final concentration of 2.5 mg/ml. Sample was deposited on glow-discharged Quantifoil R1.2/1.3 Au grids, excess of liquid was blotted by Whatman filter paper and the grids were flash frozen in liquid ethane using a Vitrobot Mark IV (TFS) device. Grids were imaged on a TFS Titan Krios G3i operated at 300 kV equipped with a Gatan K3 detector and the Bioquantum energy filter aligned at zero-loss with a slit width of 20 eV. Tomograms were recorded as dose-fractionated movies using SerialEM 3.9 (Mastronarde, 2005) with the use of dose-symmetric tilt scheme (Hagen *et al*, 2016) at a nominal magnification of 64.000 corresponding to a nominal counted pixel size of 1.35 Å. A total of 60 tomograms were recorded with the angular range of −45° to +45° with 3° increment and a defocus range of −2 to −5 um.

All the tomogram pre-processing steps were done under the TomoBEAR workflow unless otherwise stated. The processing was performed on a single node with 16 CPU cores and 2 GPUs in automated fashion without user intervention. 1982 super-resolution non-gain-corrected tilt movies in TIFF format were sorted to separate folders by tomogram index and by tilt angle within these folders. Motion correction was done with MotionCor2 1.4.4 (Zheng *et al*, 2017) with 7 by 5 patches for local motion tracking, last frame as a reference frame and Fourier cropping of the output averages by a factor of 2. Raw stacks were generated from the motion-corrected images with the *newstack* program from IMOD (Kremer *et al*, 1996). Dynamo tilt-series alignment routines were used to generate the fiducial alignment models that were imported to *Etomo* from IMOD and the alignment model was calculated, followed by generation of the aligned stacks. GCTF (Zhang, 2016) was used to estimate defocus values per tilt, and these values were input to *ctfphaseflip* (Xiong *et al*, 2009) from IMOD, followed by gold bead detection and erasion. CTF-corrected gold-erased aligned stacks were binned by a factor of 8, bin8 tomograms were reconstructed and subjected to modified Dynamo template matching, with the first template being a bagel with dimensions similar to apoferritin. For this dataset template matching was the last step completed with TomoBEAR, and subsequent subtomogram averaging was performed interactively. Cross-correlation peaks were extracted from the tomograms with a threshold of 7 standard deviations above the mean, comprising 21886 initial particle coordinates. Particle angles were randomized to get rid of the missing wedge, which resulted from the restricted search range in template matching step. The search range was restricted due to the high symmetry point group of the protein. After an initial alignment in SUSAN (https://github.com/KudryashevLab/SUSAN, manuscript in preparation), particles were recentered and the *Dynamo* table was converted to a Relion .star file with a TomoBEAR module. The project was imported to Relion4 (Zivanov *et al*, 2022), first round of 3D classification in bin4 and selection of good classes led to 21068 final particles, which were subjected to 3D Refinement followed by CTF refinement and polishing. This final 3D refinement was performed with the box size 200×200×200 voxels^3^ at counted pixel size. Pixel size was calibrated with an atomic model giving the final estimate of 1.331 Å/pix versus the nominal 1.378 Å/pix. The global resolution was estimated with 0.143 criterion to be 2.8 Å. The final map was sharpened with a B-factor of −70 Å^2^.

#### 3. Reprocessing the dataset EMPIAR-10452 of the ion channel RyR1

This is a previously processed dataset from the benchmarking of the hybrid StA data collection scheme (Sanchez *et al*, 2020). The tomograms contain a 15 e-/Å^2^ micrograph as a zero-tilt image. Tomogram pre-processing steps were done with the TomoBEAR workflow unless otherwise stated. All steps were run automatically without user intervention. 4099 non-gain-corrected tilt movies in MRC format were sorted to folders by tomogram index and by tilt angle within these folders. Motion correction was done with MotionCor2 1.4.4 (Zheng *et al*, 2017) with 7 by 5 patches for the high-dose zero-tilt image and with global motion tracking for tilted images. 100 raw stacks were generated from the motion-corrected images with the newstack program from the IMOD package (Kremer *et al*, 1996). Dynamo tilt-series alignment (https://wiki.dynamo.biozentrum.unibas.ch/w/index.php/Walkthrough on GUI based tilt series alignment) was used to generate and refine the fiducial models that were imported to ETOMO from IMOD and refined. At this point we used the StopPipeline command and the fiducial models were manually inspected in ETOMO and errors were corrected. Then TomoBEAR was restarted from the generation of the aligned stacks. GCTF (Zhang, 2016) was used to estimate defocus values per tilt, and these values were input for CTF phase flipping with IMOD’s *ctfphaseflip* (Xiong *et al*, 2009), followed by gold bead detection and erasing. CTF-corrected gold-erased aligned stacks were binned by a factor of 8, bin8 tomograms were reconstructed and subjected to template matching with the modified DynamoTM module, with the template being a resampled low-pass-filtered map of RyR1 from this dataset (EMD-10840). Cross-correlation volumes were examined and only 52 out of 81 tomograms were kept. One hundred top-hits were extracted per tomogram and subjected to a multi-reference alignment project in Dynamo (Castaño-Díez *et al*, 2012) with 3 classes. 3252 particles from the best class were selected and subsequent subtomogram averaging was performed interactively. After the initial alignment in SUSAN, particles were re-centered and the Dynamo table containing alignment parameters for particles was converted to a RELION .star file with a dedicated TomoBEAR module. The EMPIAR-10452 dataset consists of tilt-series collected with higher exposure on the untilted image, allowing for hybrid StA, however our goal here was to benchmark tomogram preprocessing in TomoBEAR and therefore we decided to proceed data as conventional StA and keep all tilts. This way errors in tilt-series alignment or tomographic reconstruction can limit the resolution. The project was imported to RELION4 (Zivanov *et al*, 2022), particles were subjected to 3D Refinement followed by CTF refinement and polishing, and then another round of 3D Refinement. Both 3D Refinements were done in bin1 with the box size of 320×320×320 voxels^3^. Importantly, upon import, tomograms description STAR file column rlnMicrographPreExposure was modified to account for the uneven dose distribution between tilts. The global resolution was estimated with 0.143 criterion to be 8.9 Å, similar to the previously obtained results with this data. Local resolution was estimated with the Relion4 implementation.

Mandatory dependencies: MATLAB R2021a/b, Dynamo 1.1.532, IMOD 4.11. Additional dependencies: CUDA-10.x/11.x, SUSAN, MotionCor2, GCTF, AreTomo.

## AVAILABILITY

TomoBEAR is open-source, available on GitHub https://github.com/KudryashevLab/TomoBEAR; the 80S ribosome dataset and corresponding input preset files are available as a tutorial; Deposited structures of Ribosomes, RyR1, Appoferitin will be referred here.

## ACKNOWLEDGEMENTS

The work is supported by the German Research Foundation (DFG, KU 3222/2-1), Sofja Kovalevskaja Award from Alexander von Humboldt Foundation to MK. MK is supported by the Heisenberg Award from the DFG (KU3222/3-1). NB was partially supported by the Josef Buchmann Family Foundation, RMS was partially supported by the starter fellowship from SFB807 (DFG). We thank the Core Facility for cryo-Electron Microscopy (CFcryoEM) of the Charité - Universitätsmedizin Berlin for support in acquisition of the data. CFcryoEM was supported by the German Research Foundation (DFG) through grant No. INST 335/588-1 FUGG and the Berlin University Alliance (BUA). The authors thank Daniel Castano-Diez, Kendra Leigh and Christoph Diebolder for useful discussions, Uljana Kravchenko, Xiaofeng Chu, Giulia Glorani for testing the developmental versions and providing feedback, Juan Castillo from the Max Planck Institute for Biophysics for the IT support at the Max Planck for Biophysics and the high-performance computing team at the MDC for supporting our operation at the Max-Cluster.

## AUTHOR CONTRIBUTIONS

NB developed and tested the code, VM and MK contributed prototype code, VM processed the RyR1 and apoferritin datasets, AY contributed to data analysis and the code, TS produced the apoferritin dataset, RMS contributed to the TomoBEAR code. MK acquired funding, supervised the project, wrote the manuscript with input from the other authors.

The authors declare no conflicts of interest.

## Notes

### Competing Interest Statement

The authors have declared no competing interest.

