## Supplementary Material for "Streamlined Structure Determination by Cryo-Electron Tomography and Subtomogram Averaging using TomoBEAR"

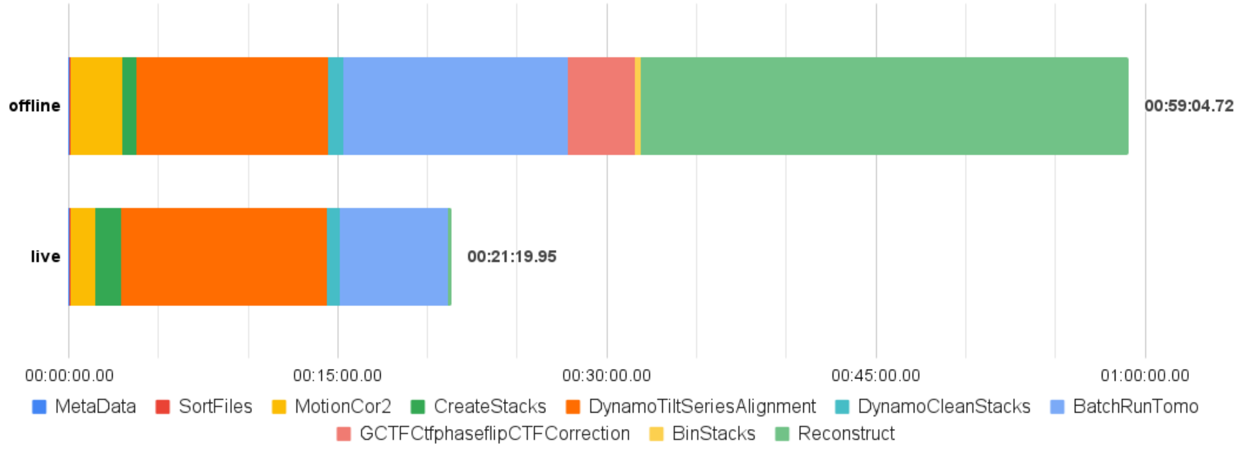


**Figure S1. Benchmarking live processing functionality with TomoBEAR.** Processing times of the HIV-1 dataset (EMPIAR-10164) for offline (conventional) and live data processing modes (hh:mm:ss.SS, where hh - hours, mm - minutes, ss - seconds, .SS - 1/100th of milliseconds).

**Supplementary Text 1: Description of Essential Modules**

The updated version can be found here:

https://github.com/KudryashevLab/TomoBEAR/wiki/Modules

**General**

The general section is not a module, but a section where all the general parameters regarding the processing and the environment can be found. Parameters which are input in this section are visible to all modules during the execution. If parameters with the same key are found in a module block, then they override parameters from the *general* section.

**MetaData**

The MetaData module collects descriptive statistics such as min, max, mean, std from the raw data.

**SortFiles**

The SortFiles module sorts the raw files into tilt-series corresponding to tomograms and links them to their corresponding folders for further processing.

**MotionCor2**

The MotionCor2 module uses MotionCor2 (Zheng *et al*, 2017) to correct for the sample movement in a given projection recorded as a dose-fractionated movie. For the options it is advised to look also into the manual of MotionCor2 as there are some more detailed descriptions.

**CreateStacks**

The CreateStacks module creates the stacks and normalizes them. There are two options for normalization, the default scheme is to divide the projections by their frame count. TomoBEAR detects automatically if you are using high-dose images from hybrid StA (Sanchez *et al*, 2020) and divides them by their corresponding frame count in contrast to low-dose images where the frame-count is different.

**DynamoTiltSeriesAlignment**

The DynamoTiltSeriesAlignment module uses the tilt stacks alignment algorithm from Dynamo (Castaño-Díez *et al*, 2012) which is the state of the art available algorithm for fiducial-based alignment. As default, reasonable parameters for many cryo-ET projects are set. Some of them are dynamically derived. The option to override non-dynamically derived parameters is available via the JSON configuration file. For troubleshooting and optimization of parameters it is possible to go to the processing folder and use the Dynamo tools as in the tutorial (https://wiki.dynamo.biozentrum.unibas.ch/w/index.php/Walkthrough_on_GUI_based_tilt_series_alignment).

**DynamoCleanStacks**

The DynamoCleanStacks module can be run after the DynamoTiltSeriesAlignment to automatically clean up the tilt stacks and to remove the projections where DynamoTiltSerieAlignment has not found gold beads. For that DynamoCleanStacks uses the output from Dynamo tilt stacks alignment which states on which projections the fiducials could be fit. The others are then removed from the tilt stacks for further processing.

**StopPipeline**

The StopPipeline module allows to stop TomoBEAR after some processing step to inspect the output and not waste computational resources if parameters need to be optimized.

**BatchRunTomo**

The BatchRunTomo is a versatile module performing IMOD operations(Mastronarde & Held, 2017). The detailed description can be found here: Tomography Guide for IMOD Version 4.11.
TomoBEAR can runs steps of batchruntomo:
0: Setup
1: Preprocessing
2: Cross-correlation alignment
3: Pre Aligned stack
4: Patch tracking, auto seeding, or RAPTOR
5: Bead tracking
6: Alignment
7: Positioning
8: Aligned stack generation
9: CTF plotting
10: 3D gold detection
11: CTF correction
12: Gold erasing after transforming fiducial model or projecting 3D model
13: 2D filtering
14: Reconstruction
14.5: Postprocessing on a/b axis reconstruction
15: Combine setup
16: Solvematch
17: Initial matchvol;
18: Autopatchfit
19: Volcombine
20: Post Processing with Trimvol
21: NAD (Nonlinear anisotropic diffusion)

**GCTFCtfphaseflipCTFCorrection**

The GCTFCtfphaseflipCTFCorrection module is detecting the defocus using gCTF (Zhang, 2016) for the tomograms which are reconstructed for template matching or particle cropping. The results can be examined in the processing folders.

**BinStacks**
The BinStacks module is used for binning the CTF corrected aligned stacks to be able to reconstruct them with the Reconstruct module.  Stacks with selected binnings will be produced.

**Reconstruct**
The Reconstruct module should be used after you binned the tilt stacks with the BinStacks module or used aligned tilt stack binning option greater than one. The module is set up by default to reconstruct binned stacks. If you otherwise want to reconstruct unbinned stacks you need to set up the Reconstruct module properly.

**DynamoImportTomograms**
The *DynamoImportTomograms* module generates a Dynamo Catalogue for you and inputs the tomograms to that catalogue. After that you can call the Dynamo Catalogue Manager (dcm) to generate the models for the tomograms or pick particles in them using the functionality of Dynamo Catalogue (Castaño-Díez *et al*, 2016).

**EMDTemplateGeneration**
The *EMDTemplateGeneration* module is used to automatically download a template which is further downscaled to match your desired template matching binning. Besides that an automated routine to generate the mask is also implemented. This module needs to be run before template matching is executed.

**TemplateGenerationFromFile**
The *TemplateGenerationFromFile* module imports a map into the TomoBEAR workflow given by a path and scales it properly if the map header contains the correct pixel size; else the pixel size can be input as a parameter through the JSON-based configuration file.

**DynamoTemplateMatching**
The *DynamoTemplateMatching* module implements basically the template matching from dynamo, but on a GPU. The GPU usage is 12-15 times faster than the conventional CPU-based template matching implementation in Dynamo (Castaño-Díez *et al*, 2012). In some multi-GPU systems the speedup is not linear to the number of GPUs.

**TemplateMatchingPostProcessing**
The *TemplateMatchingPostProcessing* module takes the cross-correlation volumes generated by the *DynamoTemplateMatching* module and extracts the coordinates from the peaks found in them until a given threshold is reached. This threshold is set by default to a value of 2.5 standard deviations.

**DynamoAlignmentProject**
The *DynamoAlignmentProject* module can be set up to generate the two common Dynamo (Castaño-Díez *et al*, 2012) subtomogram averaging projects: multiple reference alignment (MRA) project and an independent half-set based refinement project. For classification we find it useful to have one or two classes with the correct reference that was used e.g. for template

matching and some classes containing only noise ("noise traps") which will attract suboptimal particles into classes for removal.

**Live Processing (currently in the development branch)**

In the "live" mode processing is available for dose-fractionated movies in .tif or .mrc format. For this user needs to set up an input *.json* file for the TomoBEAR project in the same way as for "offline" mode and use "local_live" as the execution environment instead of "local". The user should specify a number of tilt images expected per tilt-series ("minimum_files") and listening time threshold ("listening_time_threshold_in_minutes"). The latest parameter is used as a threshold to the time passed since the latest arrived movie upon which processing of the tilt-serie will start even if not all expected movies have arrived.

**Supplementary Text 2: A tutorial for structural determination of purified ribosomes (EMPIAR 100064)**

The dataset to download and use in this tutorial you may get by the following link: https://www.ebi.ac.uk/empiar/EMPIAR-10064/. In our case we used just the `mixedCTEM` data, potentially you can additionally use the `CTEM` data to be able to pick even more particles.

After downloading the data, extract it in a folder of your choice. Note that in this case

- the data is already motion corrected
- the stacks are already assembled
- the pixel size is not in the header
- the tilt angles are not provided

`TomoBEAR` is able to incorporate this information along with the `JSON` file which describes the processing pipeline. If you have already cloned the TomoBEAR Github repository (https://github.com/KudryashevLab/TomoBEAR) to your local machine you can find in the configurations folder a file called `ribosome_empiar_10064_dynamo.json`. This file describes the processing pipeline which should be setup by `TomoBEAR` to process this data set.

The following paragraphs will explain the variables contained in the `JSON` file and the needed changes to be able to run `TomoBEAR` on your local machine. First and most importantly, you need to pass to `TomoBEAR` the path to the data and the processing folder. This must be done in the section `"general": {}` of the `JSON` file.

```
   "general": {
        "project_name": "Ribosome",
        "project_description": "Ribosome EMPIAR 10064",
        "data_path": "/path/to/the/ribosome/data/*.mrc",
        "processing_path": "/path/to/the/processing/folder/",
        "expected_symmetrie": "C1",
        "apix": 2.62,
           "tilt_angles": [-60.0, -58.0, -56.0, -54.0, -52.0, -50.0, -48.0,
-46.0, -44.0, -42.0, -40.0, -38.0, -36.0, -34.0, -32.0, -30.0, -28.0, -26.0,
-24.0, -22.0, -20.0, -18.0, -16.0, -14.0, -12.0, -10.0, -8.0, -6.0, -4.0,
-2.0, 0.0, 2.0, 4.0, 6.0, 8.0, 10.0, 12.0, 14.0, 16.0, 18.0, 20.0, 22.0,
24.0, 26.0, 28.0, 30.0, 32.0, 34.0, 36.0, 38.0, 40.0, 42.0, 44.0, 46.0, 48.0,
50.0, 52.0, 54.0, 56.0],
        "rotation_tilt_axis":-5,
        "gold_bead_size_in_nm": 9,
        "template_matching_binning": 8,
        "binnings": [2, 4, 8],
        "reconstruction_thickness": 1400,
        "as_boxes": false
  },
```

Everything else should be fine for now and the processing can be started. To run the `TomoBEAR` on the Ribosome dataset you need to type in the following command in the command window of MATLAB

```
runTomoBear("local", "/path/to/ribosome_empiar_10064_dynamo.json")
```

or if you are using a compiled version of `TomoBEAR` and have everything set up properly, type in the following command on the command line from the `TomoBEAR` folder

```
./run_tomoBEAR local /path/to/ribosome_empiar_10064_dynamo.json /path/to/defaults.json
```

When you follow all the steps thoroughly, `TomoBEAR` should run up to the first appearance of `StopPipeline`. That means the following modules will be executed. Add them to the end of .json file

```
   "MetaData": {
    },
    "CreateStacks": {
    },
    "DynamoTiltSeriesAlignment": {
    },
    "DynamoCleanStacks": {
    },
    "BatchRunTomo": {
        "skip_steps": [4],
        "ending_step": 6
    },
    "StopPipeline": {
    },
```

This can take a while, as the result of this segment `TomoBEAR` will create a folder structure with subfolders for the individual steps. You can monitor the progress of the execution in shell and by inspecting the contents of the folders. Upon success of an operation, a file `SUCCESS` is written inside each folder. If you want to rerun a step you can terminate the process, change parameters, remove the `SUCCESS` file (or the entire subfolder) and restart the process. Here the stacks have already been assembled, so neither `"Motioncorr2": {}`, not `"SortFiles": {}` modules were not needed. Here the key functionality is performed by `"DynamoTiltSeriesAlignment": {}` (a recommended tutorial: https://wiki.dynamo.biozentrum.unibas.ch/w/index.php/Walkthrough_on_GUI_based_tilt_series_alignment) after which the projections containing low number of tracked gold beads are excluded by `"DynamoCleanStacks": {}`. Finally, the output is converted into an IMOD project.
The running time depends on your infrastructure and setup. After `TomoBEAR` stops you can inspect the fiducial model in the folder of `"BatchRunTomo": {}` which you can find in your processing folder.

```
cd /path/to/your/processing/folder/5_BatchRunTomo_1
```

Now you can inspect the alignment of every tilt stack one after the other and can possibly refine it if needed. For that you can use the following command. Please replace `xxx` with the tomogram number(s) that you want to inspect or * for all tomograms.

```
etomo tomogram_xxx/*.edf
```

When `etomo` starts - choose the `fine alignment` step which should be magenta-colored if everything went fine for that tomogram and then click on `edit/view fiducial model` to start `3dmod` with the right options to be able to refine the gold beads. If you do not see the green circles - please go to the imod command window, Edit -> Object I-> Type-> activate scattered and increase the size of the circle to e.g. 5. To refine the gold beads click on `Go to the next big residual` in the window with the stacked buttons from top to bottom and the view in the view port window should change to the location of a gold bead with a big residual. Now see if you can centre the marker better on the gold bead with the right mouse button. It is important that you don't put it on the peak of the red arrow, but center it on the gold bead. When you are finished with this gold bead press again on the `Go to next big residual` button. After you are finished with re-centering the marker on the gold beads you need to press the `Save and run tiltalign` button.
After you finish the inspection of all the alignments in all tomograms you can start `TomoBEAR` again as previously and it will continue from where it stopped up to the next `StopPipeline` section. To continue running `TomoBEAR` on the Ribosome data set you need to type in as previously the following command in the command window of MATLAB
`runTomoBear("local", "/path/to/ribosome_empiar_10064_dynamo.json")`
or if you are using a compiled version of `TomoBEAR` and have everything set up properly type in the following command on the command line from the `TomoBEAR` folder

```
./run_tomoBEAR local /path/to/ribosome_empiar_10064_dynamo.json /path/to/defaults.json
```

`TomoBEAR` should now detect that it has stopped at the previous step `StopPipeline` and continue from where it stopped. The following excerpt from the `ribosome_empiar_10064_dynamo.json` file is describing what `TomoBEAR` needs to do next.

```
   "BatchRunTomo": {
        "starting_step": 8,
        "ending_step": 8
    },
    "GCTFCtfphaseflipCTFCorrection": {
    },
    "BatchRunTomo": {
        "starting_step": 10,
        "ending_step": 13
```

```
    },
    "BinStacks": {
    },
    "Reconstruct": {
    },
    "DynamoImportTomograms": {
    },
    "EMDTemplateGeneration": {
        "template_emd_number": "3420",
        "flip_handedness": true
    },
    "DynamoTemplateMatching": {
    },
    "TemplateMatchingPostProcessing": {
        "cc_std": 2.5
    },
```

This segment performs estimation of defocus to calculate the Contrast Transfer Function (CTF) using GCTF and subsequent CTF-correction using Ctfphaseflip from IMOD (`"GCTFCtfphaseflipCTFCorrection": {}`). You can inspect the quality of fitting by going into the folder `8_GCTFCtfphaseflipCTFCorrection_1` and typing

```
imod tomogram_xxx/slices/*.ctf
```

and making sure that the Thon rings match the estimation. If not - play with the parameters of the `GCTFCtfphaseflipCTFCorrection` module.

Then binned aligned CTF-corrected stacks are produced by `"BinStacks": {}` and tomographic reconstructions are generated for the binnings specified in the section `"general": {}`. In this example the particles are picked using template matching. First a template from EMDB is produced at a proper voxel size, then `"DynamoTemplateMatching": {}` creates cross-correlation (CC) volumes which can be inspected. Finally, highest cross-correlation peaks, over 2.5 standard deviations above the mean value in the cross-correlation volume are selected for extraction to 3D particle files, the initial coordinates are stored in the `particles_table` folder as a file in the dynamo table format. You can inspect the cross correlation volumes and set the threshold for more or less stringent extraction of particles.

In the section below you will find subtomogram classification projects that should produce you a reasonable structure. They first use multi-reference alignment projects with a true class and so-called "noise trap" classes to first classify out false-positive particles produced by template matching, this happens at binning which was used for template matching. In the end of the segment you should have a reasonable set of particles in the best class.

```
   "DynamoAlignmentProject": {
        "iterations": 3,
```

```
        "classes": 4,
        "use_noise_classes": true,
        "use_symmetrie": false
    },
    "DynamoAlignmentProject": {
        "iterations": 3,
        "classes": 4,
        "use_noise_classes": true,
        "use_symmetrie": false,
        "selected_classes": [1]
    },
    "DynamoAlignmentProject": {
        "iterations": 3,
        "classes": 4,
        "use_noise_classes": true,
        "use_symmetrie": false,
        "selected_classes": [1]
    },
    "DynamoAlignmentProject": {
        "iterations": 3,
        "classes": 4,
        "use_noise_classes": true,
        "use_symmetrie": false,
        "selected_classes": [1]
    },
    "DynamoAlignmentProject": {
        "iterations": 3,
        "classes": 4,
        "use_noise_classes": true,
        "use_symmetrie": false,
        "selected_classes": [1]
    },
    "DynamoAlignmentProject": {
        "iterations": 3,
        "classes": 4,
        "use_noise_classes": true,
        "use_symmetrie": false,
        "selected_classes": [1]
    },
    "DynamoAlignmentProject": {
        "iterations": 3,
        "classes": 3,
        "use_noise_classes": true,
```

```
        "use_symmetrie": false,
        "selected_classes": [1],
        "box_size": 1.10,
        "binning": 4
    },
    "StopPipeline": {
    },
```

After subtomogram classification projects are done, you should have a reasonable set of particles in the best class which you should select. To select the best class you need to go into the last `DynamoAlignmentProject` folder before the last produced `StopPipeline` folder, and then go to `alignment_project_1_bin_y/mraProject_bin_y/results/iteQQQQ/averages/` (where `iteQQQQ` corresponds to the pre-last iteration folder) and type

```
imod average_ref_CCC_ite_QQQQ.em
```

to open produced average for each class `CCC` to identify the best class to use further.
Variables binning `y`, pre-last iteration number `QQQQ` and class numbers `CCC` can depend on parameters used in `DynamoAlignmentProject`. But if you repeat instructions provided in this tutorial, this should be the folder

```
23_DynamoAlignmentProject_1/alignment_project_1_bin_4/mraProject_bin_4/result
s/ite0012/averages/
```

where you may find files `average_ref_001_ite_0012.em`, `average_ref_002_ite_0012.em`, and `average_ref_003_ite_0012.em` corresponding to the produced averages for 3 classes, from which you should choose the best one.
Once you have selected the best class, insert corresponding class number in the list `[]` as a value of the parameter `"selected_classes"` to the following section to be executed by `TomoBEAR`:

```
   "DynamoAlignmentProject": {
        "classes": 1,
        "iterations": 1,
        "use_noise_classes": false,
        "swap_particles": false,
        "use_symmetrie": false,
        "selected_classes": [3],
        "binning": 4,
        "threshold":0.8
    },
```

The section above is called a single reference project, which will split the particles of the previously selected best class into two equally sized classes (called even/odd halves) with subsequent alignment of the particles in those halves to produce corresponding averages. This division will be needed further when unbinned data will be produced to be able to calculate the resolution of the resulting averaged map using Fourier Shell Correlation (FSC) curve.
After the first single reference project introduced above you will need to process tomograms by similar projects but at lower binnings in order to reduce the voxel size up to unbinned data to get the information corresponding to the highest possible resolution to be achieved using the current dataset. At this point automated workflow is finished as the user needs to play with the masks, particle sets, etc.
You may want to try to use the following example of the end section of `JSON` file in order to have experience of processing tomograms at lower binnings to produce unbinned data to finally be able to calculate resolution of your ribosome density map as a result of the first experience with `TomoBEAR`

```
"DynamoAlignmentProject": {
     "classes": 1,
     "iterations": 1,
     "use_noise_classes": false,
     "swap_particles": false,
     "use_symmetrie": false,
     "selected_classes": [1,2],
     "binning": 2,
     "threshold":0.9
 },
 "BinStacks":{
     "binnings": [1],
     "use_ctf_corrected_aligned_stack": false,
     "run_ctf_phaseflip": true
 },
 "Reconstruct": {
     "reconstruct": "unbinned"
 },
 "DynamoAlignmentProject": {
     "classes": 1,
     "iterations": 1,
     "use_noise_classes": false,
     "swap_particles": false,
     "use_symmetrie": false,
     "selected_classes": [1,2],
     "binning": 1,
     "threshold":1
 }
```

Consider, that after performing first single reference project you need to select both halves from the previous step by setting `"selected_classes": [1,2]` (in order to keep all particles) while producing one class by setting `"classes": 1` at the subsequent steps of binning reduction using `"DynamoAlignmentProject": {}` module.

If you get `out of memory` error while running some of `"DynamoAlignmentProject": {}` at lower binnings (especially the last one), you may put additional parameter `"dt_crop_in_memory": 0` to the corresponding `"DynamoAlignmentProject": {}` sections in order to prevent keeping the whole tomogram in memory for processing. For example, in this tutorial the size of the one of unbinned tomograms is ~72Gb, while for binning 2 it is near 9Gb.
Finally, to estimate resolution of produced by `TomoBEAR` results, you need to use the following `Dynamo` command in `MATLAB`:

```
fsc = dfsc(path_to_half1, path_to_half2, 'apix', 2.62, 'mask', path_to_mask,
'show', 'on')
```

where `path_to_half1` and `path_to_half2` are paths to the pre-last iteration results of the last `DynamoAlignmentProject` folder, which in this tutorial are located in `29_DynamoAlignmentProject_1/alignment_project_1_bin_1/mraProject_bin_1_eo/results/ite0006/averages`, where you may find files `average_ref_001_ite_0006.em` and `average_ref_002_ite_0006.em` corresponding to the averages made from halves of the resulting particles set. You also need to use a mask to filter averages for FSC calculation, and the accuracy of the used mask has an impact on the resolution estimation. Appropriate mask to use for the initial resolution estimation you may find in the last `DynamoAlignmentProject` folder in a file called `mask.em` (in this tutorial `path_to_mask` is `29_DynamoAlignmentProject_1/mask.em`).

After that you should get a similar FSC curve to the following one:

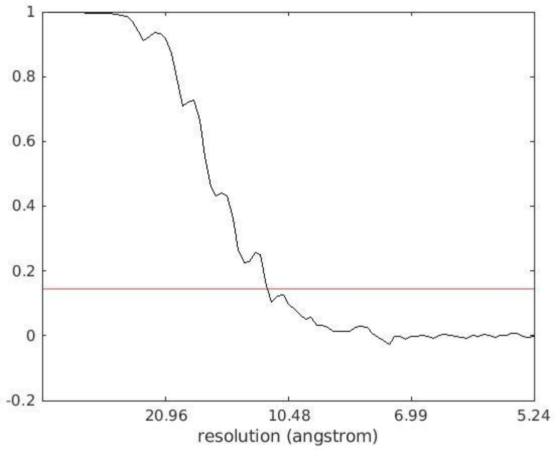


where in Red we added the `FSC = 0.143` threshold to estimate the global resolution of the final map, which in our case for the final set of 4003 ribosome particles reached 11.3 Å.
